# Deep sea sediments associated with cold seeps are a subsurface reservoir of viral diversity

**DOI:** 10.1101/2020.09.08.284018

**Authors:** Zexin Li, Donald Pan, Guangshan Wei, Weiling Pi, Jiang-Hai Wang, Yongyi Peng, Lu Zhang, Yong Wang, Casey R.J. Hubert, Xiyang Dong

## Abstract

In marine ecosystems, viruses exert control on the composition and metabolism of microbial communities, thus influencing overall biogeochemical cycling. Deep sea sediments associated with cold seeps are known to host taxonomically diverse microbial communities, but little is known about viruses infecting these microorganisms. Here, we probed metagenomes from seven geographically diverse cold seeps across global oceans, to assess viral diversity, virus-host interaction, and virus-encoded auxiliary metabolic genes (AMGs). Gene-sharing network comparisons with viruses inhabiting other ecosystems reveal that cold seep sediments harbour considerable unexplored viral diversity. Most cold seep viruses display high degrees of endemism with seep fluid flux being one of the main drivers of viral community composition. *In silico* predictions linked 14.2% of the viruses to microbial host populations, with many belonging to poorly understood candidate bacterial and archaeal phyla. Lysis was predicted to be a predominant viral lifestyle based on lineage-specific virus/host abundance ratios. Metabolic predictions of prokaryotic host genomes and viral AMGs suggest that viruses influence microbial hydrocarbon biodegradation at cold seeps, as well as other carbon, sulfur and nitrogen cycling via virus-induced mortality and/or metabolic augmentation. Overall, these findings reveal the global diversity and biogeography of cold seep viruses and indicate how viruses may manipulate seep microbial ecology and biogeochemistry.

## Introduction

Marine cold seeps are typically found at the edges of continental shelves and feature mainly of gaseous and liquid hydrocarbons from deep geologic sources^1, 2^. Seep fluids may come from thermogenic oil and gas systems that have been present for long periods of time in lower strata, indicating underlying oil and gas reservoirs^3, 4^. In the context of global climate change, methane and other short-chain alkanes escaping from deep sea cold seep sediments can reach the atmosphere, exacerbating the greenhouse effect^5^. Understanding the biogeochemical cycling in marine sediments associated with cold seeps is thus important for meeting critical energy and climate challenges.

Cold seeps are a chemosynthetic ecosystem and contain an extensive diversity of archaea and bacteria which play important roles in hydrocarbon metabolism^6, 7^. These microbial populations are not only highly active in influencing seep biogeochemistry at the sediment-water interface^8^, but also contribute to a variety of biological processes such as sulfate reduction, sulfur oxidation, denitrification, metal reduction and methanogenesis within the seabed^2, 8^. Viruses have also been observed in cold seep sediments. Epifluorescence microscopy of sediments from the Gulf of Mexico revealed that viral-like particle counts and virus-to-prokaryote ratios at cold seeps were significantly higher than in surrounding sediments, suggesting these habitats may be hot spots for viruses^9^. This agrees with elevated microbial activity at cold seeps driven by the availability of energy-rich substrates supplied from below. In addition, novel viruses have also been discovered in methane seep sediments^10^. These findings suggest that cold seeps harbour abundant and undiscovered viruses potentially influencing their microbial hosts and consequently, biogeochemical cycling at cold seeps.

Knowledge of the ecological roles of viruses in deep sea sediments has been limited by difficulties in sampling and extracting viral particles (virions) from sediments^11^. In recent years, developments in sequencing and bioinformatics have enabled the analysis of viruses recovered from metagenomes sequenced without prior virion separation. These methods have greatly advanced viral ecology from the identification of novel viruses to the global distribution of viruses. Studies from a variety of environments such as thawing permafrost^12^, mangroves^13^, arctic lakes^14^, freshwater lakes^15^, and particularly seawater^16–18^ have suggested that prokaryotic viruses act as key agents in natural ecosystems via a range of interactions with their microbial hosts. Viruses can influence organic carbon and nutrient turnover by top-down control of microbial abundance via lysis of cells and the subsequent release of cellular contents during lytic infection^19^. They can also reprogram host metabolism through horizontal gene transfer, or via auxiliary metabolic genes (AMGs) in their genomes that are expressed during infection. In peatland soils along a permafrost thaw gradient in Sweden, virus-encoded glycoside hydrolases were found to play a role in complex carbon degradation^12^. In freshwater lakes fed with sediment-derived methane, some viruses were found to encode subunits of particulate methane monooxygenase, suggesting that they may augment bacterial aerobic methane oxidation during infection^20^. In recent years, studies are starting to reveal the presence and abundance of viruses in deep sea sediments^11, 21, 22^, thus deep sea sediments associated with cold seeps present a unique opportunity to study viruses and their interactions with hosts in a chemosynthetic ecosystem often dominated by anaerobic methane oxidation. Several metagenomic sequencing efforts have been undertaken on cold seep sediments^2^ such that the extracted DNA includes genomes of viruses in these sediments, yet most studies have focused exclusively on genomes bacteria and archaea, neglecting the viruses.

In this study, we sought to expand the understanding of viral diversity and the ecological role of viruses in deep sea sediments associated with cold seeps. To this end, 28 publicly available marine sediment metagenomes from seven cold seeps around the world were analyzed to recover genomes of viruses in cold seep communities. Characterizing the diversity of these viral communities enabled predictions about the organisms they may be infecting and identification of AMGs potentially mediating ecological roles of viruses in these habitats. Our findings reveal the global diversity and biogeography of seep viruses and their role in benthic microbial ecology and biogeochemistry.

## Methods

### Collection of metagenomic datasets for deep sea cold seeps

Metagenomic data sets were compiled from 28 sediment samples collected from seven cold seep sites across the global oceans (**Figure 1)**. These sites were: Haakon Mosby mud volcano (HM); Eastern North Pacific ODP site 1244 (ENP); Mediterranean Sea, Amon mud volcano (MS); Santa Monica Mounds (SMM); Eastern Gulf of Mexico (EGM); Scotian Basin (SB); and Western Gulf of Mexico (WGM) (**Supplementary Table 1 and references therein**). Except for EGM and SB, metagenomic datasets along with metadata were downloaded from NCBI Sequence Read Archive and NCBI BioSample databases (https://www.ncbi.nlm.nih.gov). Sample collection and DNA sequencing of samples from EGM and SB are described in detail elsewhere^23, 24^.

**Figure 1.**
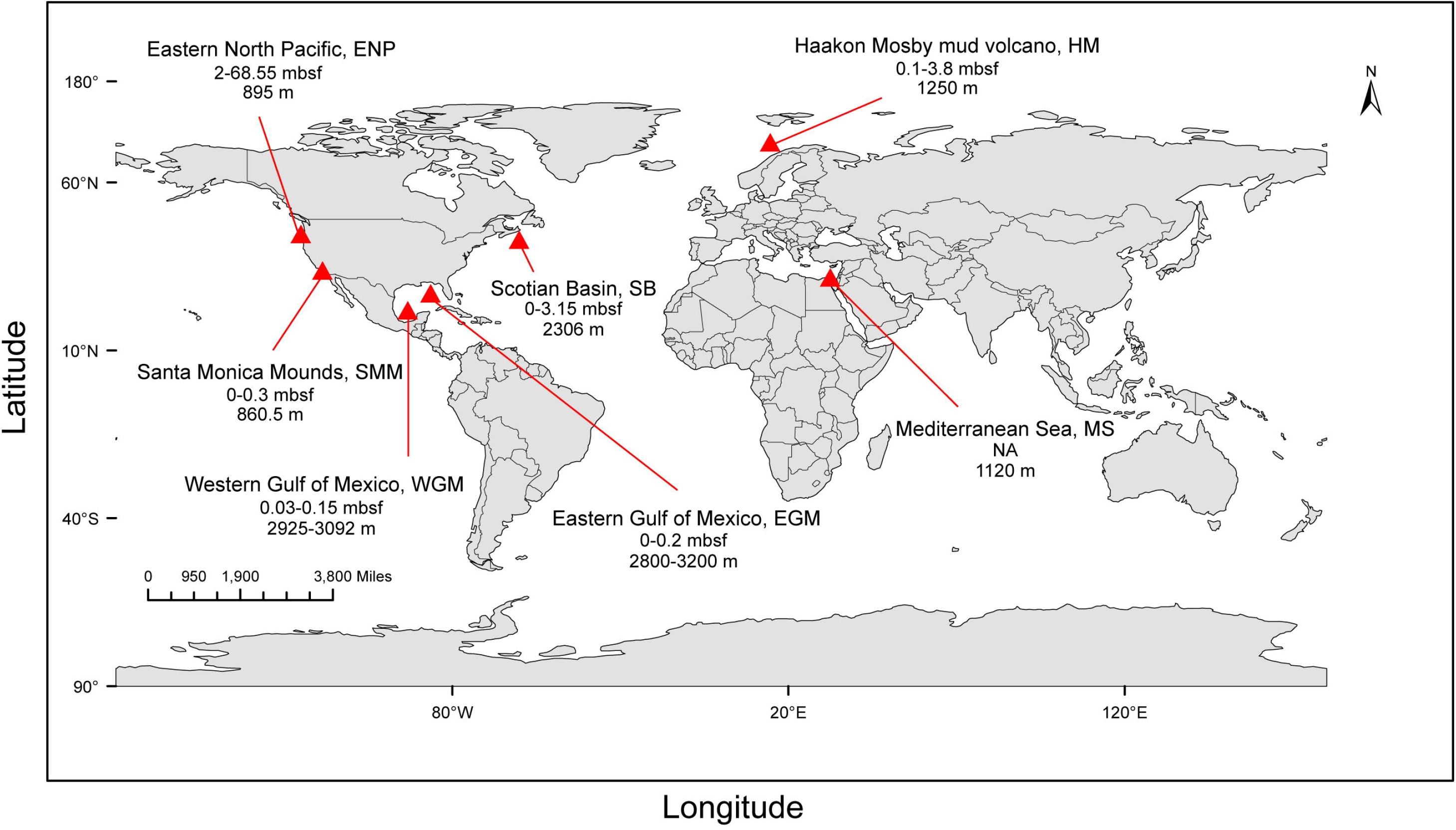
Geographic distribution of sampling sites where metagenomic data were collected. Locations of cold seep sites indicating the site name and abbreviation, sampling depth range in meters below seafloor (mbsf) and water depth in meters below sea level (m).

### Taxonomic profiling of microbial communities

To explore the prokaryotic composition of each sample, 16S rRNA gene fragments (i.e. miTags) were extracted from metagenomic raw reads using the phyloFlash pipeline^25^. Extracted 16S miTags were mapped to the SILVA SSU rRNA reference database (v132)^26^ and assigned an approximate taxonomic affiliation (nearest taxonomic unit, NTU).

### Metagenomic assembly

Raw reads were quality-controlled by trimming primers and adapters and filtering out artifacts and low-quality reads using the Read_QC module within the metaWRAP pipeline^27^. Quality-controlled reads from each metagenome were individually assembled using MEGAHIT v1.1.3^28^ (default parameters). Short contigs (<1000 bp) were removed.

### Generation of prokaryotic metagenome-assembled genomes

For each assembly, contigs were binned using the binning module (parameters: --maxbin2 --metabat1 --metabat2) and consolidated into a final bin set using the Bin_refinement module (parameters: -c 50 -× 10) within metaWRAP^27^. All the produced bin sets were aggregated and dereplicated at 95% average nucleotide identity (ANI) using dRep v2.3.2 (parameters: -comp 50 -con 10 -sa 0.95)^29^, resulting in a total of 592 species-level metagenome-assembled genomes (MAGs). Taxonomy of each MAG was initially assigned using GTDB-Tk v0.3.3^30^ based on the Genome Taxonomy Database (GTDB, http://gtdb.ecogenomic.org) taxonomy R04-RS89^31^. The results were further refined using maximum-likelihood phylogeny inferred from a concatenation of 120 bacterial or 122 archaeal marker genes produced by GTDB-Tk. Bacterial and archaeal trees were built using RAxML v8^32^ called as follows: raxmlHPC-HYBRID -f a -n result -s input -c 25 -N 100 -p 12345 -m PROTCATLG -x 12345. Genomes were finally classified using the naming system of the NCBI taxonomy^33^.

### Identification of viral contigs

Viral contigs were recovered from metagenome assemblies using VirSorter v1.0.5^34^ and VirFinder v1.1^35^. Only contigs ≥10 kb were retained, based on the following criteria^17^: (1) VirSorter categories 1, 2, 4 and 5; (2) VirFinder score ≥0.9 and *p*<0.05; (3) both VirSorter categories 1-6 and VirFinder score ≥0.7 and *p*<0.05. The identified contigs from each assembly were then compiled and clustered at 95% nucleotide identity using CD-HIT v4.8.1 (parameters: -c 0.95 -d 400 -T 20 -M 20000 -n 5)^36^, producing 2885 viral OTUs (vOTUs). These may represent a mixture of free viruses, proviruses and/or actively infecting viruses^12^. Completeness of viral genomes was estimated using the CheckV pipeline^37^. CheckV and VIBRANT v1.2.1^38^ were used to infer temperate lifestyles by identifying viral contigs that contain provirus integration sites or integrase genes.

### Comparisons to viral sequences from other environments by protein clustering

To place the 2885 vOTUs in broader context, they were compared to viral contigs in public databases: (i) GOV 2.0 seawater^17^ (n=195728); (ii) wetland sediment^39^ (n=1212); (iii) Stordalen thawing permafrost^12^ (n=1896). For each viral contig, open reading frames (ORFs) were called using Prodigal v2.6.3^40^ and the predicted protein sequences were used as input for vConTACT2^41^. We followed the protocol published in protocols.io (https://www.protocols.io/view/applying-vcontact-to-viral-sequences-and-visualizi-x5xfq7n) for the application of vConTACT2 and visualization of the gene-sharing network in Cytoscape v3.7.2^42^ (edge-weighted spring-embedded model). Viral RefSeq (v85) was selected as the reference database, and Diamond was used for the protein-protein similarity method. Other parameters were set as default.

### Viral taxonomic assignments

To identify the taxonomic affiliations of the vOTUs, ORFs predicated from Prodigal v2.6.3 were aligned against the viral NCBI Viral RefSeq V94 using BLASTp (E-value of <0.0001, bitscore ≥50)^13, 17, 43^. The BLASTp output was then imported into MEGAN v6.17.0 using the Lowest Common Ancestor (LCA) algorithm for taxonomic analysis^44^.

### Abundance profiles

RPKM (Reads per kilobase per million mapped reads) values were used to represent relative abundances of viruses and microorganisms. To calculate the RPKM values of each viral contig or MAG, quality-controlled reads from each sample were mapped to a viral contig database or to contigs compiled from the 592 MAGs with BamM v1.7.3 ‘make’ (https://github.com/Ecogenomics/BamM). Low quality mappings were removed with BamM v1.7.3 ‘filter’ (parameters: --percentage_id 0.95 --percentage_aln 0.75). Filtered bam files were then passed to CoverM v0.3.1 (https://github.com/wwood/CoverM) to generate coverage profiles across samples (parameters: contig mode for viral contigs, genome mode for MAGs, --trim-min 0.10 -- trim-max 0.90 --min-read-percent-identity 0.95 --min-read-aligned-percent 0.75 -m rpkm).

### Virus-host prediction

Four different *in silico* methods^12, 39, 45^ were used to predict virus-host interactions. *(1) Nucleotide sequence homology*. Sequences of vOTUs and prokaryotic MAGs were compared using BLASTn. Match criteria were ≥75% coverage over the length of the viral contig, ≥70% minimum nucleotide identity, ≥50 bit score, and ≤0.001 e-value. *(2) Oligonucleotide frequency (ONF)*. VirHostMatcher v1.0^46^ was run with default parameters, with d ^*^ values ≤0.2 being considered as a match. *(3) Transfer RNA (tRNA) match*. Identification of tRNAs from prokaryotic MAGs and vOTUs was performed with ARAGORN v1.265 using the ‘-t’ option^47^. Match requirements were ≥90% length identity in ≥90% of the sequence by BLASTn^18^. *(4) CRISPR spacer match*. CRISPR arrays were assembled from quality-controlled reads using crass v1.0.1 with default parameters^48^. CRISPR spacers were then matched against viral contigs with ≤1 mismatch over the complete length of the spacer using BLASTn. For each matching CRISPR spacer, the repeat from the same assembled CRISPR array was compared against the prokaryotic MAGs using BLASTn with the same parameters, creating a virus-host link. Among potential linkages, *cas* genes of putative microbial hosts were inspected further using MetaErg v1.2.2^49^. Only hits with adjacent *cas* genes were regarded as highly confident signals.

Whenever multiple hosts for a vOTU were predicted, the virus-host linkage supported by multiple approaches was chosen. Otherwise, virus-host linkage determination used a previously-reported^12^ priority order of: (1) CRISPR spacer match with adjacent *cas* gene; (2) CRISPR spacer match without adjacent *cas* gene; (3) tRNA match or nucleotide sequence homology; (4) ONF comparison

### Functional annotations of MAGs

Each MAG was first annotated using MetaErg v1.2.2^49^. The predicted amino sequences were then used as query for identification of key metabolic markers via METABOLIC v2.0^50^. For phylogenetic analysis of McrA and DsrA, amino acid sequences were aligned using the MUSCLE algorithm^51^ included in MEGA X^52^. All positions with less than 95% site coverage were eliminated. The maximum-likelihood phylogenetic tree was constructed in MEGA X using the JTT matrix-based model. The tree was bootstrapped with 50 replicates and midpoint-rooted.

### Identification of auxiliary metabolic genes

AMGs were identified based on KEGG, Pfam and VOG annotations using a combination of VIBRANT v1.2.1 and METABOLIC v2.0^50^ with default parameters. Manual inspection was used to remove non-AMG annotations. CAZyme (carbohydrate-active enzyme) genes were identified on the dbCAN2^53^ web server based on the recognition of the CAZyme signature domain found by at least two out of three tree tools (HMMER + DIAMOND + Hotpep).

### Statistical analyses

All statistical analyses were performed in R version 3.6.3. Alpha and beta diversity of viral communities were calculated using vegan package v2.5-6^54^. Shapiro-Wilk and Bartlett’s tests were employed to test the data normality and homoscedasticity prior to other statistical analysis. For beta diversity of viral communities, non-metric multidimensional scaling (NMDS) was used to reduce dimensionality using the function capscale with no constraints applied. NMDS was based on Bray-Curtis dissimilarities generated from OTU tables with viral abundances (RPKM) using the vegdist function (method ‘‘bray’’). The grouping of cold seep sites into different types^2^ (mineral-prone vs mud-prone) was individually verified using Analysis of similarity (ANOSIM). For comparison between cold seep sites, Shannon index was compared using Analysis of Variance (ANOVA) while Simpson and Chao1 indices were compared using a Kruskal-Wallis nonparametric test. For comparison of cold seep systems, Shannon index was compared using Student’s T-test while Simpson and Chao1 indices were compared using Wilcoxon signed-rank test. Pearson correlations were performed using the cor function.

### Results and Discussion

To investigate the diversity and ecological function of viruses inhabiting cold seep sediments, a 0.38 Tbp compilation of metagenomic data was recruited from public databases and analysed (**Supplementary Table 1**). Metagenomes were sequenced from 28 sediment samples obtained at seven seabed cold seeps across the global oceans, encompassing gas hydrates, oil and gas seeps, mud volcanoes and asphalt volcanoes (**Figure 1**).

### Overview of bacterial and archaeal communities

To assess the overall microbial community structure in these sediments, 16S miTags were extracted from metagenomic reads for taxonomic profiling^25^. Classification of 16S miTags at the phylum level (class level for Proteobacteria) revealed dominant bacterial lineages to be Chloroflexi (on average 23% of bacterial 16S miTags from 28 samples), Atribacteria (23%), *Gammaproteobacteria* (9%), *Deltaproteobacteria* (9%), and Planctomycetes (6%) (**Supplementary Figure 1**). In shallow sediments (<0.2 meters below the sea floor; mbsf), *Gammaproteobacteria* and *Deltaproteobacteria* were present in higher relative abundance, whereas Atribacteria and Chloroflexi predominated in deeper sediments that made up the majority of the sample set. For archaeal lineages, members from *Methanomicrobia* (phylum Euryarchaeota) were on average 30% of archaeal miTags, followed by Bathyarchaeota (TACK group) at 18%, and Lokiarchaeota (Asgard group) at 16% (**Supplementary Figure 2**).

Assembly and binning of metagenomes resulted in 592 high-- or medium-quality^55^ microbial MAGs clustering at 95% ANI, nominally representing species-level groups^56^. These 460 bacterial and 132 archaeal MAGs spanned 46 known and four unclassified phyla (**Figure 2a** and **Supplementary Table 2**). Within the domain Bacteria, members of Chloroflexi (n=119 MAGs), *Deltaproteobacteria* (n=67) and Planctomycetes (n=44) were highly represented. Within the domain Archaea, MAGs were mainly affiliated with *Methanomicrobia* (n=41), Bathyarchaeota (n=21) and Lokiarchaeota (n=18). Based on the read coverage of MAGs among the samples, no single MAG was found to be present in all seven cold seeps (**Supplementary Table 3**). All seven regions harboured MAGs belonging to *Deltaproteobacteria* (n=67), Planctomycetes (n=44), WOR-3 (n=15), Bacteroidetes (n=14), Heimdallarchaeota (n=12) and Atribacteria (n=11).

**Figure 2.**
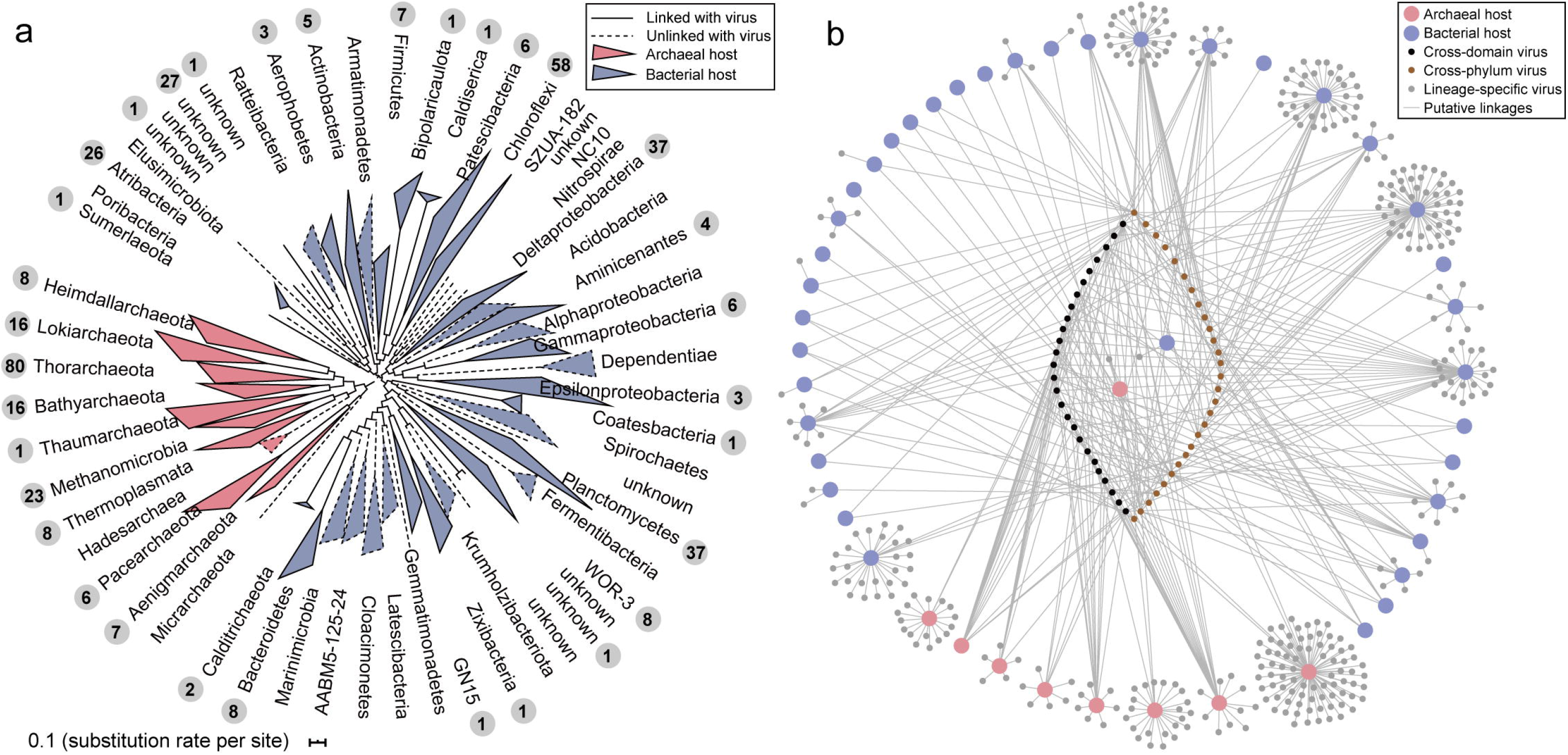
Cold seep virus-host linkages. (a) Maximum-likelihood phylogenetic tree of bacterial and archaeal MAGs at the phylum level (class level for Proteobacteria and Euryarchaeota), inferred from a concatenated alignment of 120 bacterial or 122 archaeal single-copy marker genes. Clades outlined by solid lines represent lineages predicted to include a host for one or more viral OTUs (number of vOTUs predicted to have a host within a clade is shown in grey circles). (b) Network of putative virus-host linkages. Edges indicate putative virus-host pairs. Large nodes represent bacterial (blue) or archaeal (pink) hosts. Small nodes represent vOTUs coloured according to host ranges: grey, host-specific infection at or below phylum level; brown, cross-phylum infection; black, cross-domain infection.

### Viruses from cold seep sediments are diverse and novel

From the 28 bulk shotgun metagenomes, 39154 putative viral sequences were obtained, manually filtered and then clustered at 95% ANI to represent approximately species-level taxonomy^17, 57^. This gave rise to 2885 non-redundant cold seep vOTUs, each represented by contigs ≥10kb in size, including four that were ≥200 kb (**Supplementary Table 4**) possibly corresponding to huge viruses^58^. Completeness of metagenome-assembled viral genomes or genome fragments was estimated using CheckV^37^, giving rise to four different quality tiers: complete genomes (10.3% vTOUs), high-quality (4.3%), medium-quality (11.3%), and low-quality (57.7%), with the remainder 16.4% being undetermined (**Supplementary Figure 3**).

Cold seep vOTUs were abundant across all sediment samples (**Supplementary Table 5**), however a large majority (84%) of vOTUs were only present within a single cold seep site. Further analysis of viral distribution across the seven cold seep sites (ANOSIM, *R*=0.802, *p*=0.0001) also shows that cold seep viruses display a high degree of endemism, similar to what was found previously in methane seep prokaryotic communities^59^. Viral Shannon diversity, Simpson diversity and Chao1 richness were all observed to be significantly different (*p*<0.05) between the seven sites (**Supplementary Table 6**). To arrange samples into environmentally meaningful groups, the seven cold seeps were designated as mineral-prone or mud-prone systems according to their fluid flow regime^2^. Low-flux, mineral-prone systems have longer geologic history with slower emission of fluids, e.g., gas hydrates (i.e. ENP, SMM and SB) and oil and gas seeps (i.e. EGM) whereas younger mud-prone systems are high-flux, such as mud volcanoes (i.e. HM and MS), asphalt volcanoes (i.e. WGM), brine pools and brine basins. Non-metric multidimensional scaling (NMDS) analysis revealed clear dissimilarity between viral communities in mineral-prone and mud-prone systems (ANOSIM, *R*=0.558, *p*<0.001; **Figure 3a**). For the most part, viral communities from mineral-prone systems clustered together, however SB_0 (surface sediment from 0.0 mbsf) deviated from other Scotian Basin viral communities as well as those from other mineral-prone seeps. Other factors thus also contribute to the structuring of the viral community, possibly including sediment depth (**Figure 3a**). Shannon diversity, Simpson diversity and Chao1 richness of viral communities were significantly higher in mineral-prone than in mud-prone seep systems (**Figure 3b**). Overall these results suggest that fluid flux is an important driver of viral community compositions in cold seep sediments.

**Figure 3.**
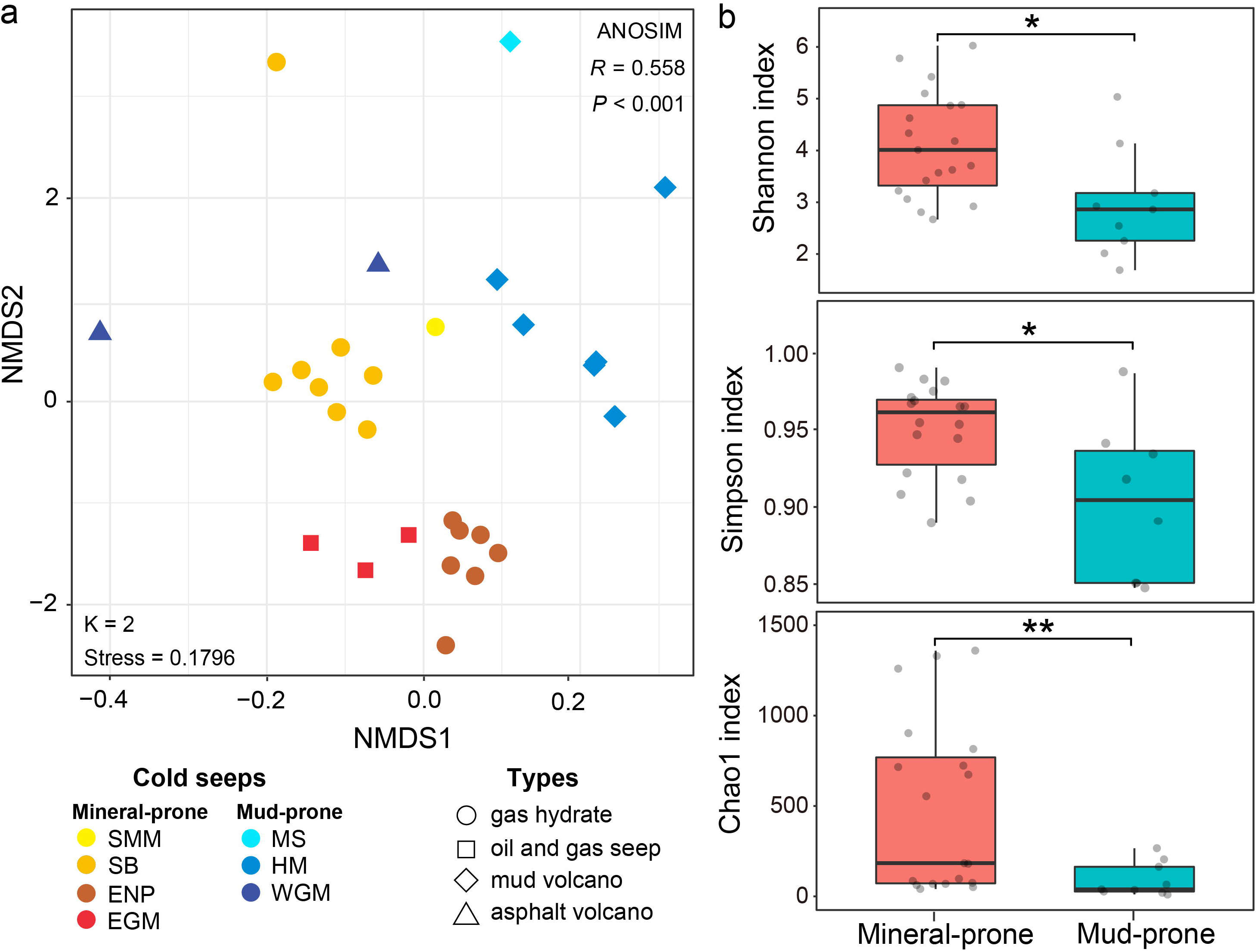
Comparison of viral community diversity between mineral-prone and mud-prone cold seeps. (a) NMDS analysis of a Bray-Curtis dissimilarity matrix calculated from RPKM values of vOTUs. ANOSIM was applied to test for the difference between viral communities in mineral-prone (n=19) or mud-prone (n=9) systems. (b) Shannon, Simpson and Chao1 indices of the viral community diversity from mineral-prone and mud-prone cold seeps. Asterisks denote significance, with * indicating *p*<0.05, and ** indicating *p*<0.01.

To investigate the relationship between cold seep vOTUs and publicly available virus sequences from a broader diversity of ecosystems, a gene-sharing network was constructed using vConTACT2^41^. Such a weighted network can assign sequences into viral clusters (VCs) at approximately the genus level. Cold seep sediments, seawater, wetland and permafrost vOTUs were grouped into 3082 VCs (**Figure 4a** and **Supplementary Table 7**). Only 17 VCs were shared amongst all ecosystems (**Figure 4b**). The limited extent of clustering between viral genomes sampled from the various ecosystems may reflect a high degree of habitat specificity for viruses. Among cold seep sediment viruses, 1742 out of 2885 vOTUs were clustered into 804 VCs, with the majority (78.7%) not encountered in any other ecosystem. This suggests that most cold seep viruses may be endemic to cold seeps (**Figure 4b** and **Supplementary Table 8**). Among the 2885 cold seep vOTUs, only 162 clustered with wetland-derived vOTUs, 154 with seawater-derived vOTUs, and 95 with permafrost-derived vOTUs (**Supplementary Table 8**). Very few cold seep viral vOTUs (~0.7%) clustered with taxonomically known genomes from Viral RefSeq (**Figure 4a**), which is a much lower proportion compared to viruses recently discovered in soils using a similar approach^60^. Similarly, attempted taxonomic assignment of cold seep vOTUs using whole genome comparisons against 2616 known bacterial and archaeal viruses from NCBI RefSeq (version 94) left >96% unclassified. The remainder were assigned to the *Caudovirales* order, specifically *Podoviradae* (n=35), *Myoviradae* (n=34) and *Siphoviradae* (n=27) (**Figure 4c**). These analyses show that cold seep sediments harbour considerable unexplored viral diversity.

**Figure 4.**
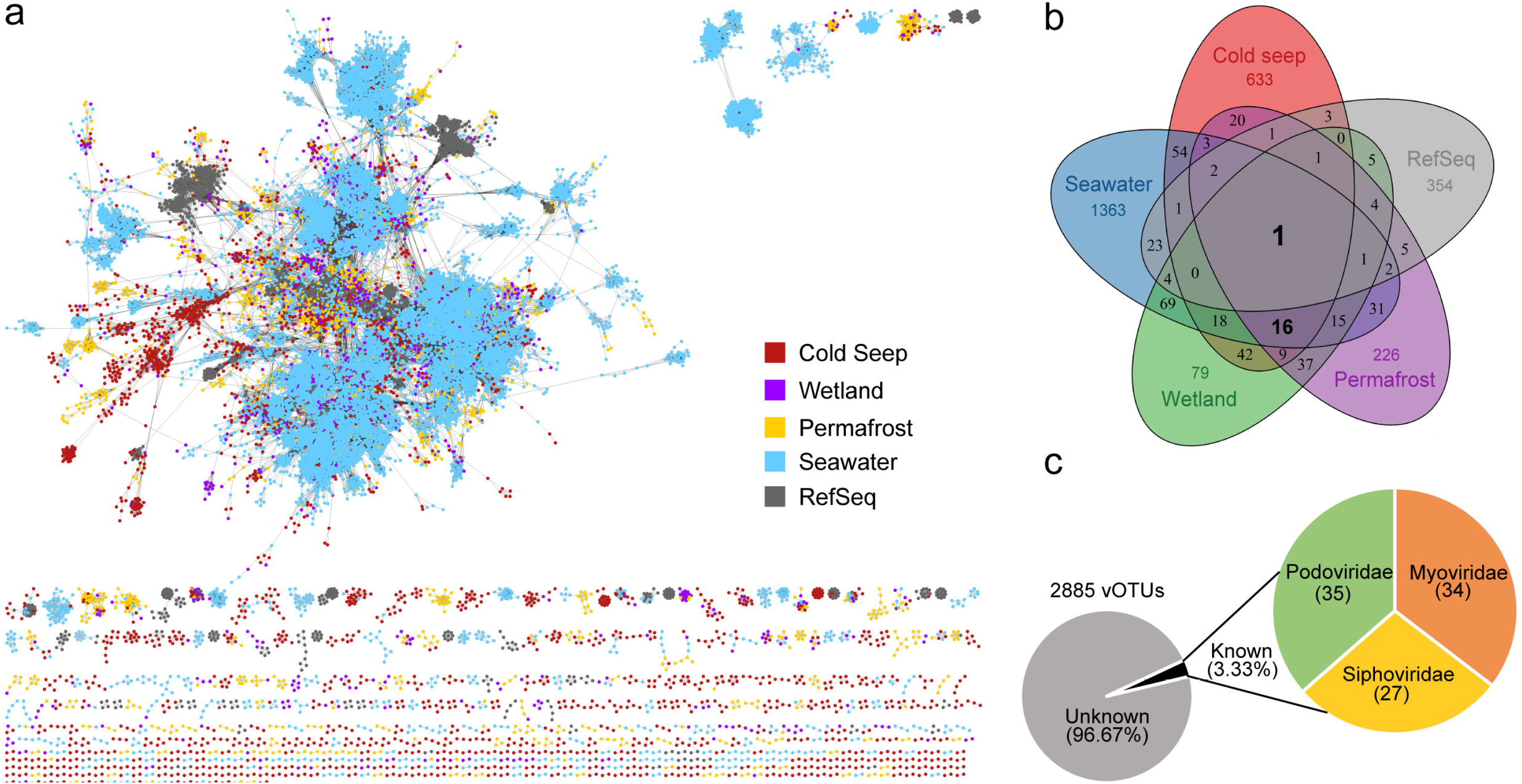
Taxonomic diversity of cold seep viruses. (a) Gene-sharing network of viral sequence space based on assembled viral genomes from cold seep sediment, wetland, permafrost, seawater and RefSeq prokaryotic viral genomes. Nodes represent viral genomes and edges indicate similarity based on shared protein clusters. (b) Venn diagram of shared viral clusters among the four environmental virus data sets and RefSeq. (c) Taxonomic assignments of vOTUs.

### Viral lifestyles, virus-host linkages and host-linked viral abundance

Comparing sequence similarity, oligonucleotide frequencies, tRNA sequences and CRISPR-spacers^61^, putative hosts were predicted for 14.2% of the 2885 cold seep vOTUs (**Supplementary Table 9**). Consistent with previous observations^61, 62^, most of these vOTUs are predicted to have narrow host ranges, with only 54 vOTUs potentially exhibiting a broader host range across several phyla. 26 vOTUs were linked to both bacterial and archaeal hosts, suggesting existence of viral infection across domains (**Figure 2b**). To minimise the impact of potential false positives, 203 low-confidence host predictions were excluded from the analysis. For virus-host pairs with the greatest confidence, predicted prokaryotic hosts spanned 9 archaeal and 23 bacterial phyla, with the most frequent predictions being Thorarchaeota (19% of virus-host pairs) and Chloroflexi (14%) (**Figure 2a**). A considerable proportion (40%) of cold seep vOTUs were linked to archaea, including members of Bathyarchaeota, the Asgard group, *Methanomicrobia*, Thaumarchaeota and *Thermoplasmata*. Such broad ranges for archaeal viruses have not been reported previously in natural systems^12, 61^. Based on the presence of functional marker genes within MAGs, predicted hosts included two aerobic methanotrophic *Methylococcales* (**Supplementary Table 10**), 13 anaerobic methane-oxidizing archaea (e.g. ANME-1 and ANME-2, **Supplementary Figure 5a**), one non-methane multi-carbon alkane oxidizer within *Methanosarcinales* (**Supplementary Figure 5a**), 16 sulfate reducers mostly belonging to *Deltaproteobacteria* (**Supplementary Figure 5b**), and numerous respiring and fermentative heterotrophs (**Supplementary Table 10**). The genome of the sulfate reducer *Desulfobacterales* 8_GM_sbin_oily_21 also harboured genes possibly encoding akyl-/arylalkylsuccinate synthases related to anaerobic degradation of longer alkanes and aromatic hydrocarbons. These results suggest that viruses may influence the carbon and sulfur cycling via the lysis of populations mediating biogeochemical processes in cold seeps, where sulfate reduction is coupled to the anaerobic oxidation of methane and other seeping hydrocarbons. Predicted hosts were also identified within the candidate phyla radiation (six vOTUs are predicted to infect Patescibacteria) and DPANN archaea (13 vOTUs are predicted to infect Pacearchaeota or Aenigmarchaeota). Due to limited metabolic capabilities and small cell sizes, many CPR and DPANN organisms are likely to be obligate symbionts of other bacteria and archaea^63^. The impact of viral infection on obligate symbionts and any consequences for the larger organisms hosting those symbionts are not yet known, although it has been suggested that they may protect those hosts from viral predation^63^.

Based on abundances determined by read mapping, targeted hosts were predicted for >20% of the cold seep viral community (**Figure 5a**). When grouped at the phylum level (class level for Proteobacteria and Euryarchaeota), the composition of predicted microbial hosts agreed well with that of their viruses (**Figure 5b**). This is supported by regression modelling of the abundances of hosts and lineage-specific viruses (**Figure 5c**). By applying metagenomic read recruitment, most viruses have higher genome coverage compared to their hosts, suggesting that most taxa may be undergoing active viral replication and possibly lysis at the time of sample collection^64^. Lineage-specific virus/host abundance ratios (i.e. VHR) for most taxa were greater than one with Thorarchaeota being the highest at 10^2.5^ (**Figure 5d**), indicating a high level of active viral genome replication. This is in accordance with the presence of higher abundances of viral particles detected by epifluorescence microscopy in cold seep sediments compared to non-cold seep sediments in the Gulf of Mexico^9^. Thus in cold seep sediments, viral lysis may be a major top-down factor^65^, contributing to significant microbial mortality. In addition, based on their contigs containing integrase genes and/or being located within their host genomes, at least 372 cold seep vOTUs were predicted to be lysogenic (i.e. temperate viruses, **Supplementary Figure 4** and **Supplementary Table 4**).

**Figure 5.**
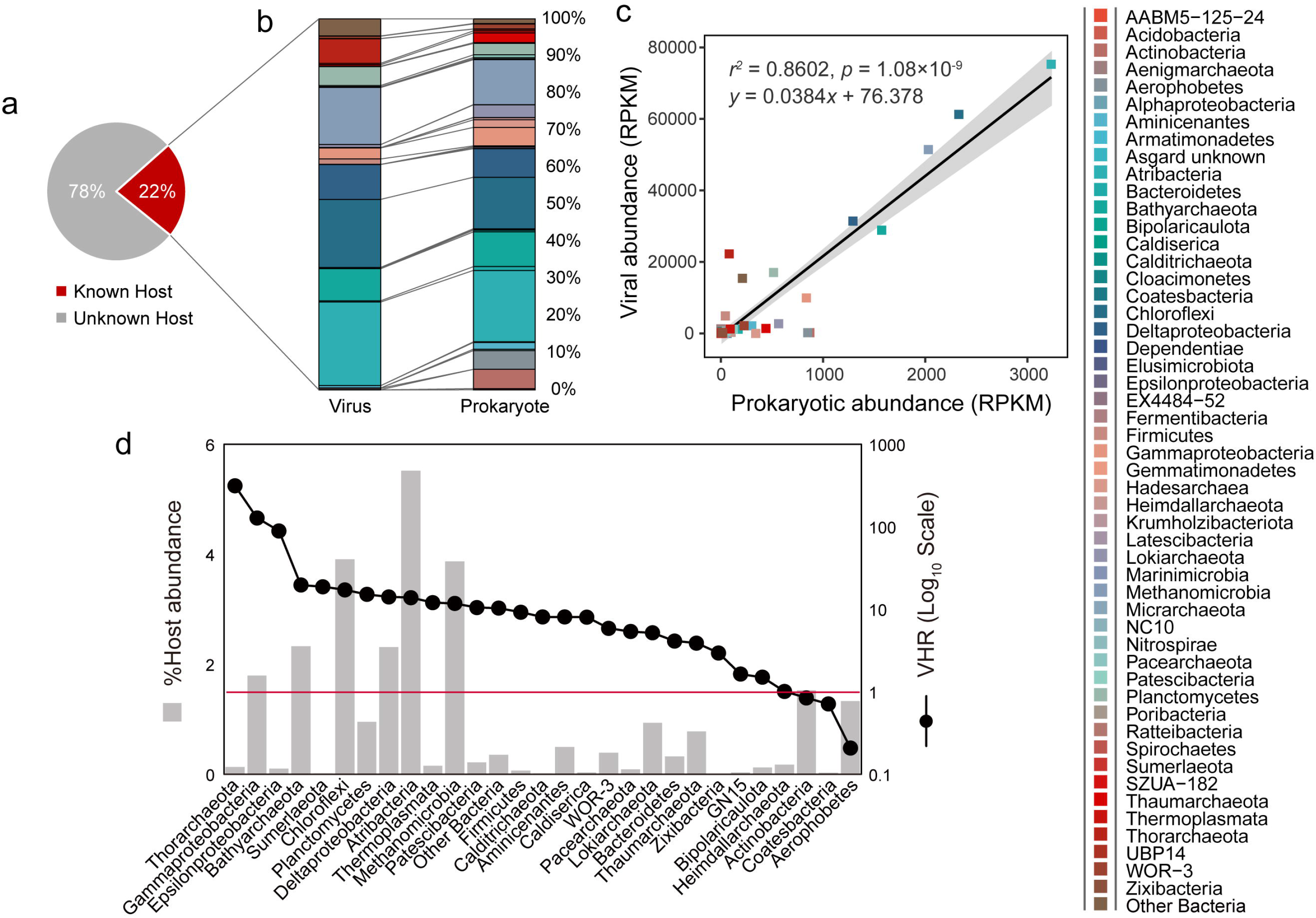
Relative abundance patterns of viruses and their predicted hosts in cold seep sediments. (a) Percentage of vOTUs based on relative abundance in which a host was predicted or not. (b) Relative abundances of vOTUs and their predicted hosts grouped by the host taxonomy (c) Significant Pearson correlation between relative abundances of viruses and their hosts (calculated by normalized mean coverage depth, reads per kilobase mapped reads: RPKM). (d) Lineage-specific virus-host abundance ratios (VHR) of for all predicted microbial hosts. The red line indicates a 1:1 ratio. Predicted hosts in (b) and (c) are in different colours as shown in the colour bar on the right side.

### Viral AMGs involved in carbon, sulfur and nitrogen transformations

To further understand how viruses might affect the biogeochemistry of cold seep sediments, viral contigs encoding AMGs that supplement host metabolism during infection were examined. Overall, cold seep viruses tended to encode AMGs for cofactor/vitamin and carbohydrate metabolism. A significant portion also encoded AMGs for amino acid and glycan metabolism (**Figure 6a**). We identified 70 genes encoding carbohydrate-active enzymes (CAZymes), related to the initial breakdown of complex carbohydrates, with 22 of them affiliated to glycoside hydrolyases (**Figure 6b**). These 22 genes, spanning 16 glycoside hydrolase families (**Supplementary Table 11**), were predicted to function in polymer hydrolysis, typical of bacteria and/or archaea (e.g. Planctomycetes and Thorarchaeota)^66^. Two *mmoB* genes encoding soluble methane monooxygenase regulatory protein B were identified in viral contigs, which might be associated with aerobic methane oxidation^67, 68^. No other AMGs directly related to key functional genes involved in initial activation of hydrocarbons were identified. However, many genes potentially involved in downstream hydrocarbon biodegradation pathways were identified, e.g., acetate-CoA ligase (*acd*), acetyl-CoA synthetase (*acs*), acetyl-CoA decarbonylase/synthase (*cdhD* and *cdhE*), 5,6,7,8-tetrahydromethanopterin hydro-lyase (*fae*), anaerobic carbon-monoxide dehydrogenase (*cooS*), 5,10- methylenetetrahydromethanopterin reductase (*mer*), and heterodisulfide reductase subunit C2 (*hdrC2*) (**Supplementary Table 12**). These genes might aid in bacterial fermentative or respiratory consumption of metabolites produced from oxidation of hydrocarbons and other complex substrates.

**Figure 6.**
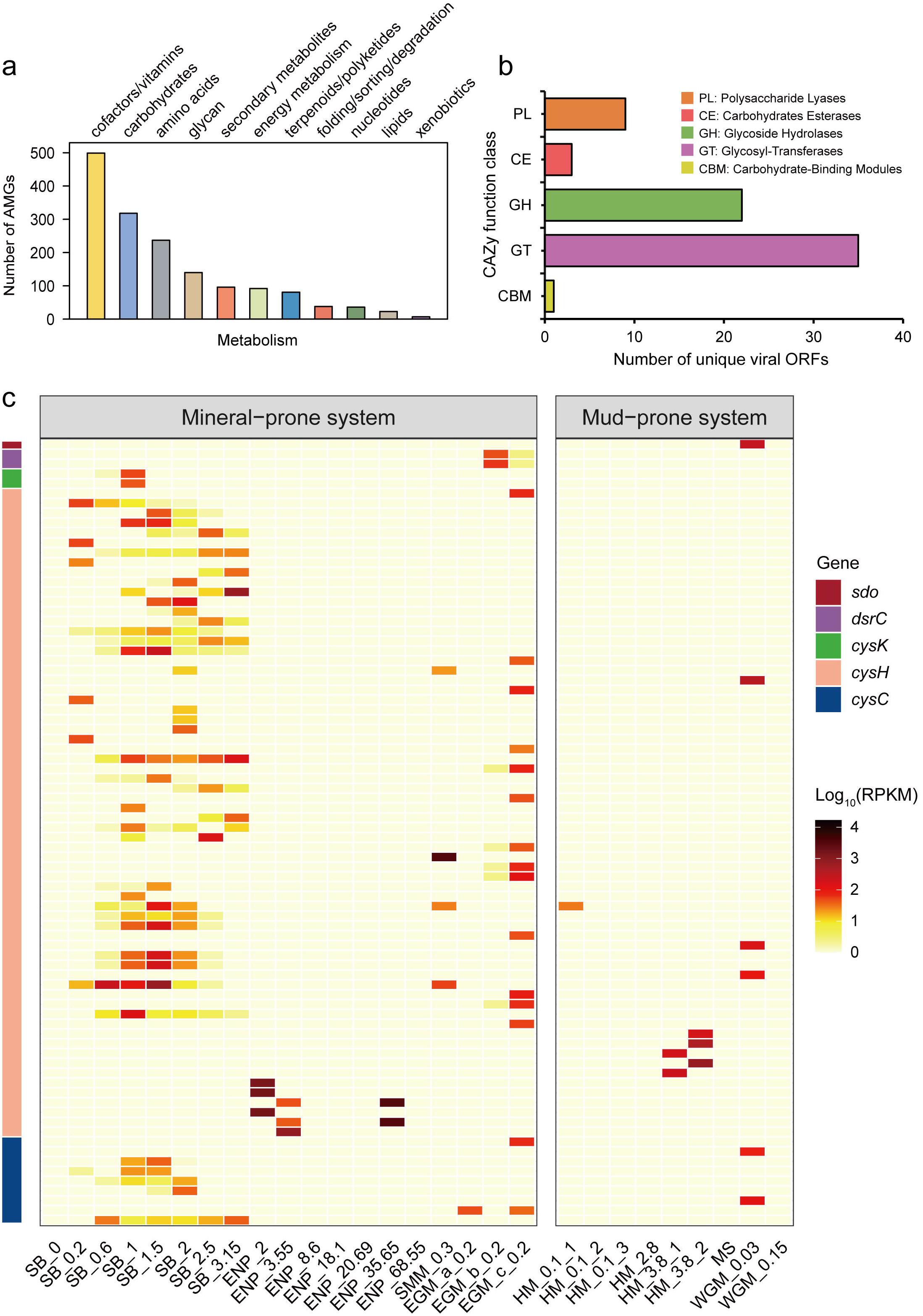
Profiles of virus-encoded auxiliary metabolic genes (AMGs). (a) Classification of AMGs into KEGG metabolic categories. (b) Classification of viral ORFs encoding carbohydrate-active enzymes. (c) Relative abundance of AMGs associated with sulfur metabolism. Gene abbreviations: sulfur dioxygenase (*sdo*), dissimilatory sulfite reductase related protein (*dsrC*), cysteine synthase (*cysK*), phosphoadenosine phosphosulfate reductase (*cysH*), adenylylsulfate kinase (*cysC*).

The most common AMG related to sulfur metabolism within the viral contigs was phosphoadenosine phosphosulfate reductase (*cysH*), predicted to participate in assimilatory sulfate reduction (**Supplementary Table 12**). Viral *cysH* has also been found in viral sequences obtained from the rumen^69^, a deep freshwater lake^15^ and sulfidic mine tailings^70^. Other related enzymes in the assimilatory sulfate reduction pathway including adenylylsulfate kinase (*cysC*) and cysteine synthase (*cysK*) were also identified but only in relatively small number of viral sequences (**Figure 6c**). These genes likely facilitate host utilization of reduced sulfur compounds during infection, providing viruses with some fitness advantage. AMGs related to sulfate assimilation were less prevalent in mud-prone systems (**Figure 6c**), possibly due to low sulfate availability in mud-prone systems, as a result of low sulfate intrusion into sediments caused by rapid rates of upward fluid flow from the subsurface^2^. Two *dsrC* genes were identified in viral contigs, and may be involved in dissimilatory sulfur metabolism^16^. One viral contig encoded a sulfur dioxygenase (*sdo*) for facilitating sulfur oxidation, with the predicted host being a *Deltaproteobacteria* (**Supplementary Tables 9 and 12**). Numerous contigs contained *nosD* (encoding a nitrous oxidase accessory protein) and two contigs contained *nrfA*, (encoding cytochrome c nitrite reductase; **Supplementary Table 12**), suggesting that viruses might also manipulate nitrogen cycling in cold seep sediments^71^.

## Conclusions

Due to the challenges of deep sea sediment sampling and laboratory cultivation of microbial communities along with their viruses, the roles that viruses play in influencing microbial mortality, ecology and evolution remains largely unexplored in marine sediments associated with cold seeps^21, 72^. In this study, in-depth exploration of untargeted *de novo* metagenomic data successfully revealed novel, abundant and diverse bacterial and archaeal viruses. Many of the putative microbial hosts for seep viruses belong to taxonomic groups with no cultured representatives. These results therefore expand the diversity of archaeal viruses, especially those infecting important archaeal lineages in hydrocarbon seep microbiomes, e.g., members of the Euryarchaeota, Bathyarchaeota, and the Asgard group. While a significant portion of the viruses appear to be lysogenic, the high read coverages for many viral genomes suggest that viral lysis is a major source of microbial mortality and biomass turnover in cold seep sediments. Virus encoded AMGs, including genes related to carbon, sulfur, and nitrogen metabolism, may augment the metabolism of prokaryotic hosts during infection, potentially altering biogeochemical processes mediated by cold seep microorganisms. As subsurface reservoirs of prokaryotic diversity and hotspots of microbial activity, cold seeps additionally represent oases of viruses and viral activity. Much remains to be revealed about the contribution of viruses to the functioning of cold seeps and other marine environments, especially with respect to their potential role in horizontal gene transfer which was not addressed in this study. With only a fraction of vOTUs identified here able to be classified, and many of them predicted to infect poorly characterized taxa, there remain large gaps in understanding the microbiology of these environments.

## Data availability

Sequences of 2885 viral contigs and 592 de-replicated metagenome-assembled genomes can be found at figshare (DOI: 10.6084/m9.figshare.12922229). All other data are available from the corresponding author upon request.

## Acknowledgements

The work was supported by the National Natural Science Foundation of China (No. 41906076), the Fundamental Research Funds for the Central Universities (No. 19lgpy90), and Guangzhou Marine Geological Survey (No. 2019C-15-229). We thank Benjamin Bolduc for help with vContact2 software, Simon Roux for help with host assignment, and Chuwen Zhang for helpful comments.

## Author contributions

X.D. designed this study. X.D., Z.L., and W.P. analyzed metagenomic data. X.D., Z.L., D.P., and G.W. interpreted data. Z.L., Y.P., and L.Z. performed viral diversity analyses. C.R.J. H. contributed part of the data. X.D., Z.L., D.P., and C.R.J. H. wrote the paper, with input from other authors.

