## Supplementary Figures 1-5 for "Deep sea sediments associated with cold seeps are a subsurface reservoir of viral diversity"

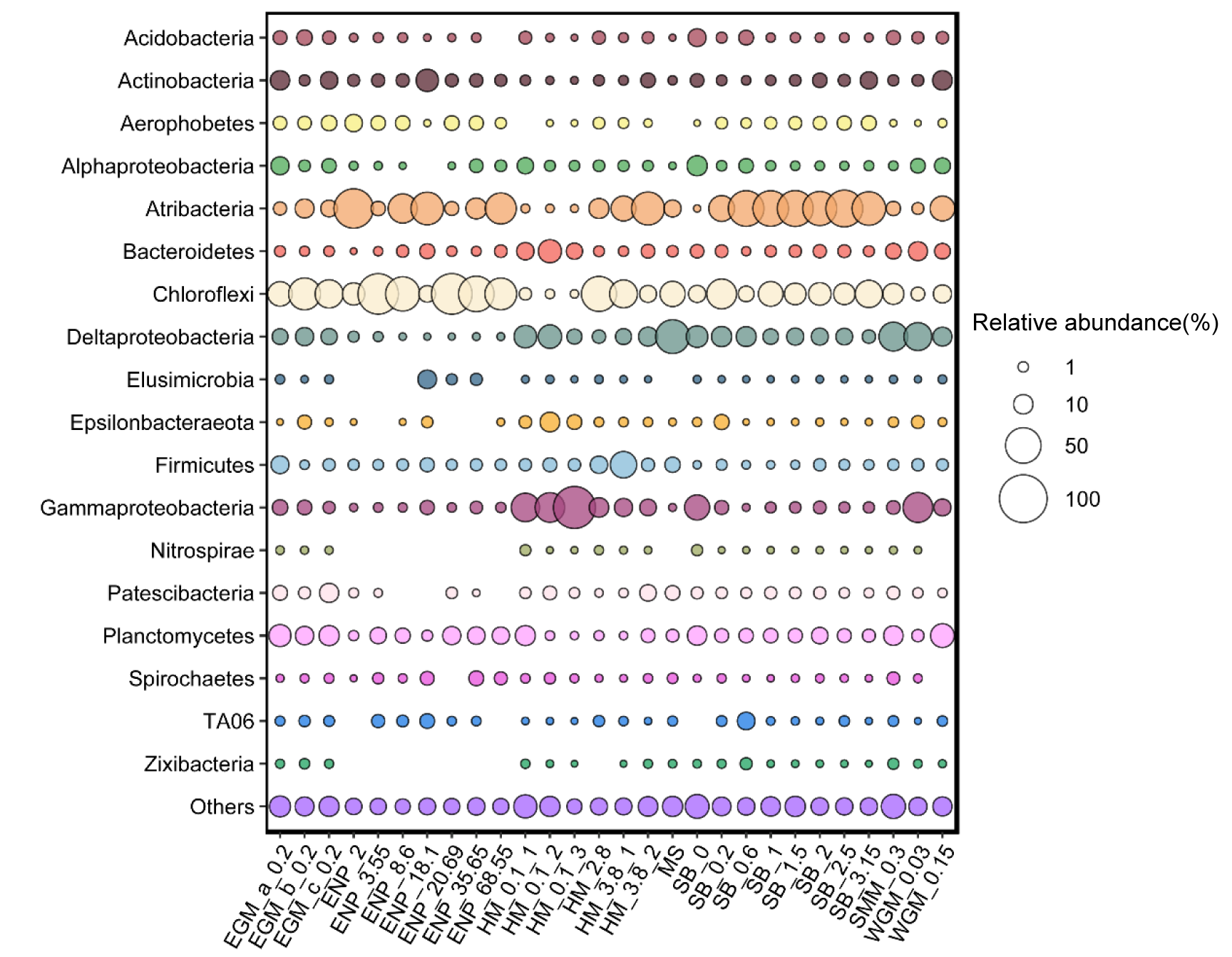

**Supplementary Figure 1.** Bubble plot of the relative abundance of bacterial 16S rRNA genes (%) in 28 cold seep sediment samples.

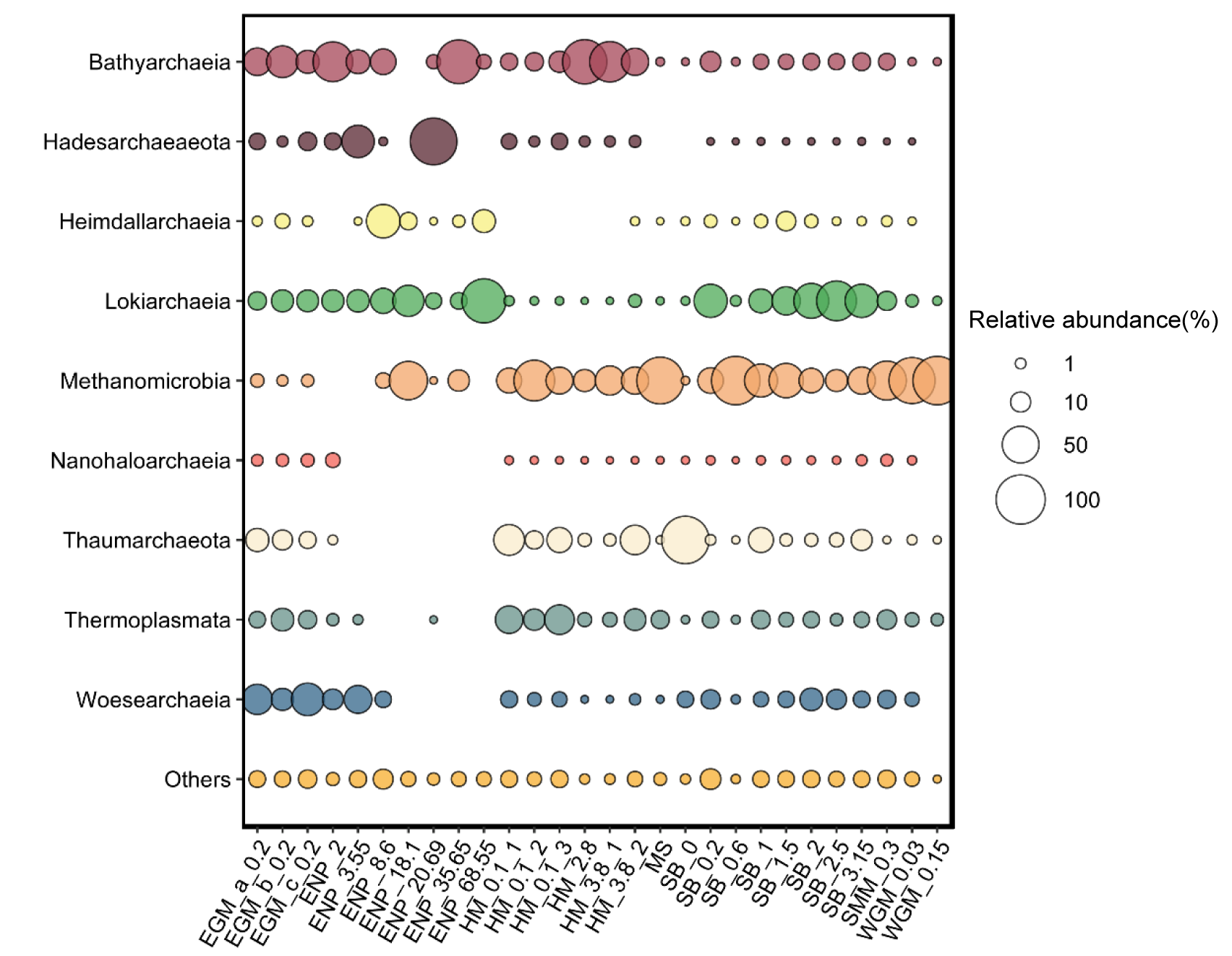

**Supplementary Figure 2.** Bubble plot of the relative abundance of archaeal 16S rRNA genes (%) in 28 cold seep sediment samples.

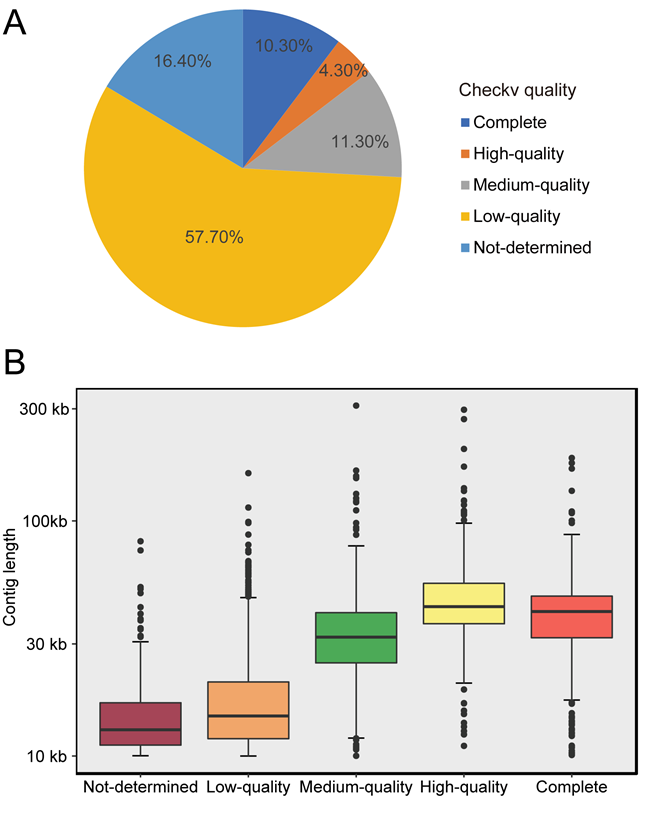

**Supplementary Figure 3.** Genome quality and completeness of vOTUs. (a) Proportion of genome quality tiers of cold seep viral populations. (b) Distribution of cold seep viral genomes across quality tiers. Quality of viral genomes were evaluated via Checkv based on estimated genome completeness. Complete: 100% completeness; high-quality: ≥ 90% completeness; medium-quality: 50-90% completeness; low-quality: < 50% completeness; not-determined: undetermined completeness.

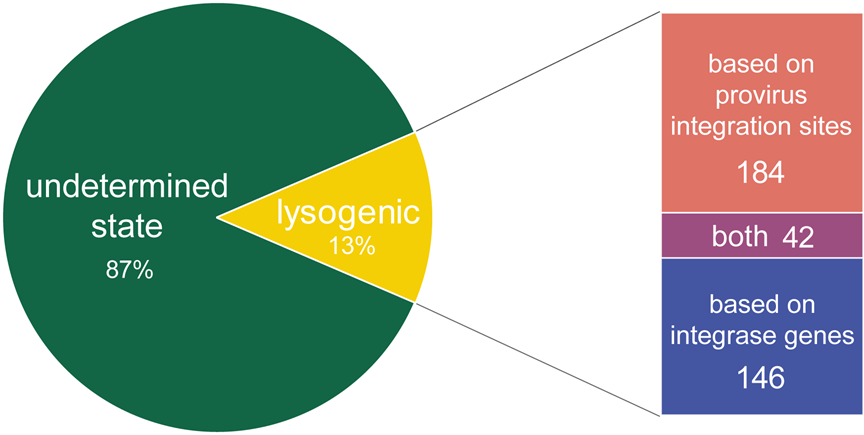

**Supplementary Figure 4.** Proportion of putatively lysogenic viruses of all the cold seep vOTUs. Lysogenic viruses were identified based on provirus integration sites and integrase genes using CheckV and VIBRANT.

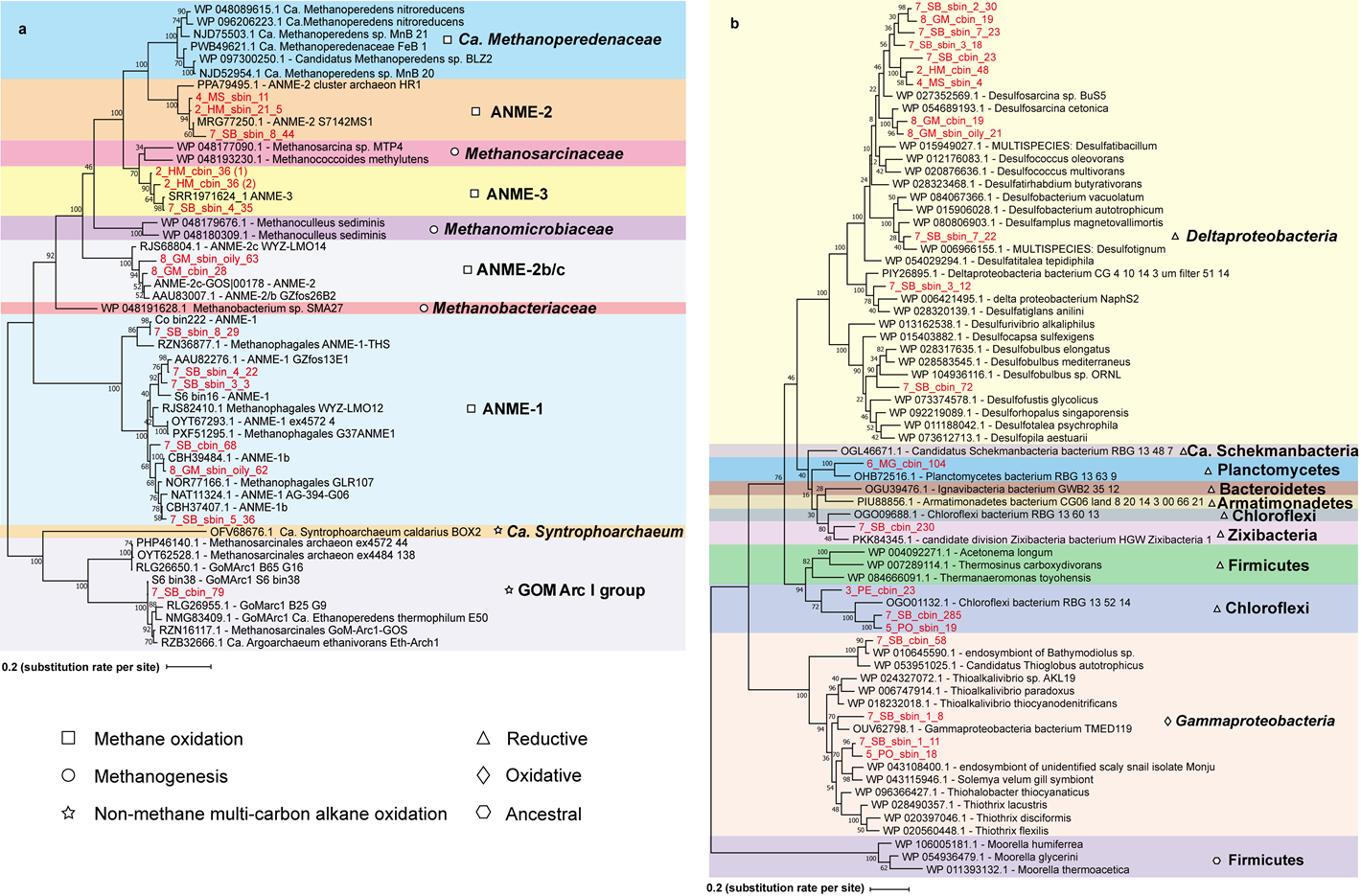

**Supplementary Figure 5.** Maximum likelihood phylogenetic trees of methyl-Coenzyme M reductase and dissimilatory sulfite reductase. Phylogenetic tree constructed based on alignments of amino acid sequences of (a) McrA and (b) DsrA genes. Bootstrap values are indicated as numbers. Genes of putative hosts in this study are highlighted in red.
